# Universal differential equations for systems biology: Current state and open problems

**DOI:** 10.1101/2024.11.29.626122

**Authors:** Maren Philipps, Nina Schmid, Jan Hasenauer

**Affiliations:** Life and Medical Sciences (LIMES) Institute and Bonn Center for Mathematical Life Sciences, Rheinische Friedrich-Wilhelms-Universität Bonn, Bonn, Germany

## Abstract

Universal Differential Equations (UDEs) combine mechanistic differential equations with data-driven artificial neural networks, forming a flexible framework for modelling complex biological systems. This hybrid approach leverages prior knowledge and data to uncover unknown processes and deliver accurate predictions. However, UDEs face challenges in efficient and reliable training due to stiff dynamics and noisy, sparse data common in biology, and in ensuring the interpretability of the parameters of the mechanistic model. We investigate these challenges and evaluate UDE performance on realistic biological scenarios, providing a systematic training pipeline. Our results demonstrate the versatility of UDEs in systems biology and reveal that noise and limited data significantly degrade performance, but regularisation can improve accuracy and interpretability. By addressing key challenges and offering practical solutions, this work advances UDE methodology and underscores its potential in tackling complex problems in systems biology.

## 1 Introduction

Systems biology aims to achieve a holistic understanding of biological processes [1]. Mathematical models have become a core tool in this endeavour, enabling researchers to represent and analyse the dynamic processes underlying biological functions. These models often take the form of differential equations that describe the temporal evolution of biological components [2]. However, as important players in biological processes and their interactions are often only partially understood, the structure of these processes remains subject to uncertainty [3, 4]. Accordingly, one of the foundational challenges in systems biology is the identification of model structures that can accurately recapitulate process dynamics solely based on experimental measurements.

Over the last decades, a broad spectrum of general-purpose methods for structure identification has been proposed [5]. Early work in this area focused on linear systems, where techniques such as state-space models were employed to infer system dynamics [6]. However, many processes are inherently nonlinear, prompting the development of more sophisticated methods [7]. These included polynomial and look-up table models, as well as neural networks and fuzzy models [5]. More recently, we have seen advancements in sparse identification methods (e.g., SINDy [8]) and the introduction of neural differential equations (NDEs) [9]. NDEs leverage the power of artificial neural networks (ANN) and offer a more flexible framework for capturing the intricate dynamics of biological systems. Yet, these data-driven methods are not designed for a flexible incorporation of prior knowledge, and the resulting models cannot be easily interpreted.

In systems biology, the exploitation of prior knowledge is often critical as the datasets are still limited as revealed by benchmarking study [10]. Furthermore, particularly for medical applications, the interpretability of the model is essential to enable decision-makers. Therefore, data-driven modelling approaches which require large datasets and provide limited interpretability are not ideally suited for systems biology. Grey box modelling approaches, which combine knowledge- and data-driven components, have been introduced to exploit prior knowledge, allow for incomplete process descriptions and facilitate model interpretation. There has been a spectrum of application-specific developments, such as the interpretable machine learning framework for perturbation biology [11], demonstrating how grey box models can effectively balance interpretability and predictive accuracy in systems biology. Furthermore, Physics-Informed Neural Networks (PINNs) [12, 13] and Universal Differential Equations (UDEs) were introduced as a generic framework [14].

Universal differential equations (UDEs) represent one of the most promising new concepts for computational modelling in systems biology, as they allow for the flexible integration of prior knowledge with data-derived terms. UDEs combine traditional differential equations with artificial neural networks (ANNs), enabling the modelling of systems where the underlying equations are partially unknown or too complex to be fully specified. The work by Rackauckas et al. has been pivotal in advancing UDEs, particularly within the Julia SciML framework, where they have been applied to a range of scientific problems, including systems biology [14]. UDEs appear to be especially powerful because they can incorporate constraints such as state positivity [15], which is essential in biological systems where certain variables (e.g., concentrations) must remain non-negative. Initial applications of UDEs in systems biology and epidemiology have demonstrated their potential to model complex biological processes with a high degree of flexibility and accuracy [14, 16]. Despite their potential, a comprehensive and unbiased assessment of the optimal strategy for training UDEs in systems biology remains unavailable.

In this study, we provide an assessment and a guide for using UDEs in systems biology. To ensure that the assessment is realistic, we address important domain-specific challenges: (1) The abundance of species as well as rate constants of biochemical processes can vary by orders of magnitude [10], and so can kinetic rates, necessitating the use of log-transformed parameters. (2) Biological systems frequently exhibit stiff dynamics [17] requiring specialised numerical solvers. (3) The measurement noise observed in systems biology often follows complex distributions, necessitating appropriate error model and maximum likelihood techniques. (4) The measurable quantities are often derived from complex combinations of biological species, formulated as observable mappings. These mappings can restrict the identifiability of model parameters, particularly when data availability is low and resolution is limited. (5) Training UDEs involves the selection of hyperparameters, such as the activation function or learning rate. (6) The flexibility of ANNs increases the susceptibility to overfitting. Moreover, UDEs need to strike a balance between the contributions of the mechanistic and ANN components. It remains unclear, how the complex ANN influences the inference of mechanistic parameters and the overall interpretability of the UDE model.

We propose methods to address these challenges by integrating best practice from machine learning and mechanistic mathematical modelling of biological processes (Fig. 1). In addition to benchmarking these methods, we evaluate UDEs for biological problems covering parts or even all these challenges. In particular, we evaluate UDE performance on synthetic problems with varying measurement noise and sparsity - spanning from simple to challenging and realistic - as well as a real-world parameter estimation problem using biological data. These case studies demonstrate the flexibility and utility of the UDE approach in addressing a range of biological modelling challenges.

**Fig. 1:**
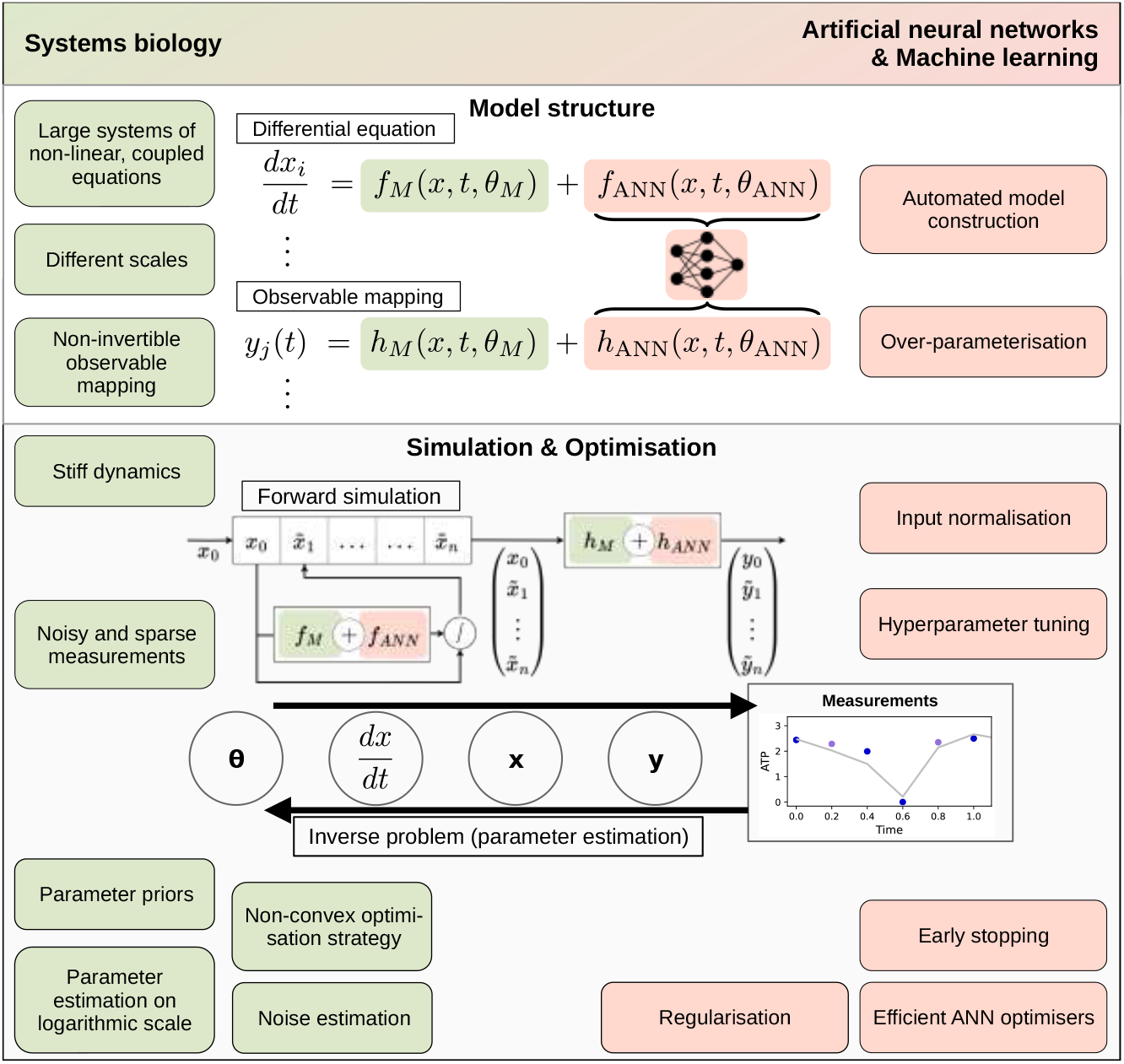
Aspects and methods from systems biology and machine learning.

Our analysis reveals that model performance and convergence deteriorate significantly with increasing noise levels or decreasing data availability, regardless of the ANN size or hyperparameter configurations. However, we identify regularisation as a key factor in restoring inference accuracy and model interpretability. To ensure reproducibility and encourage further exploration, we provide the full pipeline for model implementation, calibration, and evaluation, accessible at [GitHub link].

## 2 Results

### 2.1 Multi-start pipeline for effective UDE training

To study the challenges outlined in the introduction and to explore potential solutions, we implement a pipeline for formulating and optimising Universal Differential Equations (UDEs) (Fig. 2). The primary objective of this pipeline is to enable parameter inference for complex biological problems. Therefore, it addresses three important aspects.

**Fig. 2:**
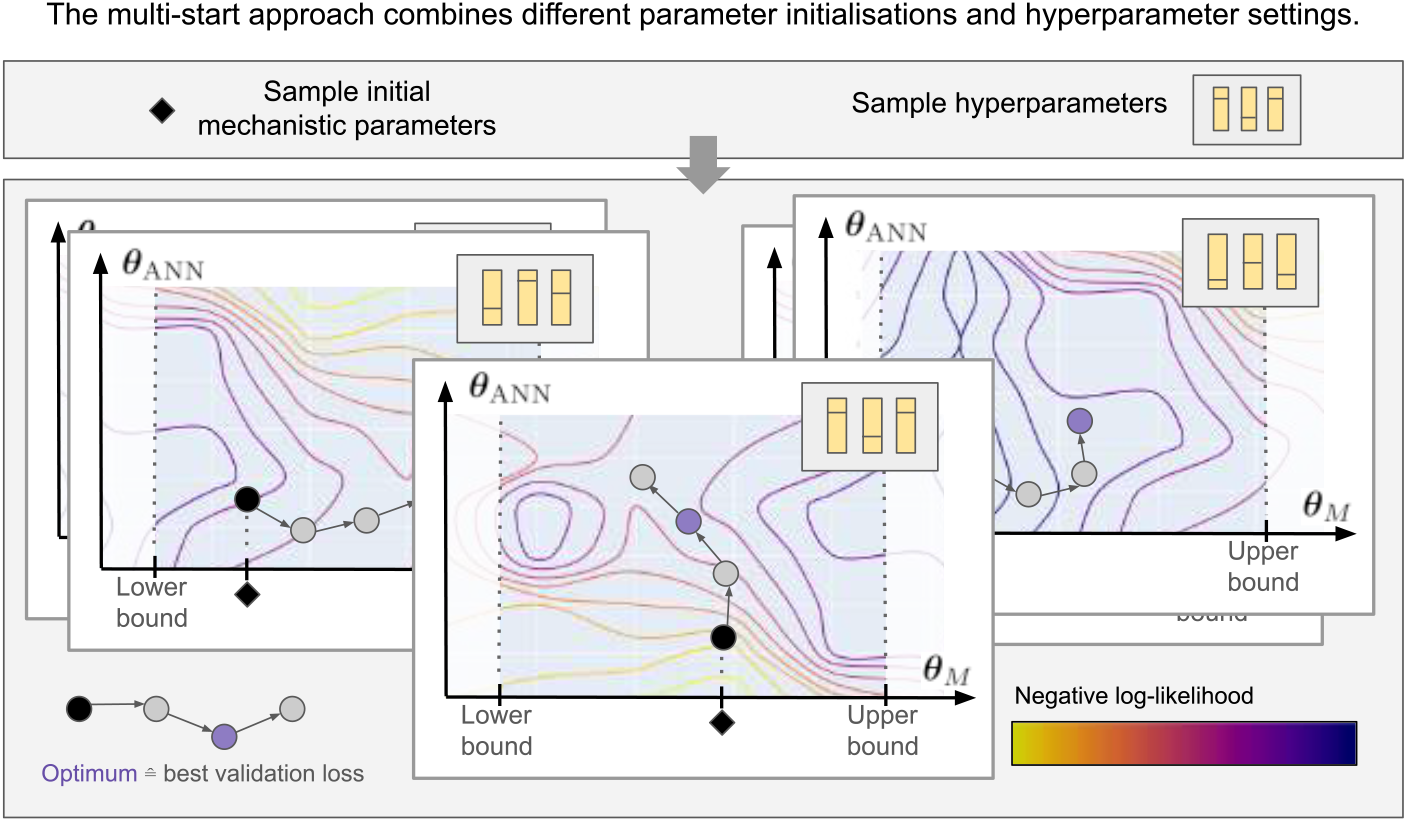
A multi-start pipeline diversifies the optimisation start points. Initial mechanistic parameters and hyperparameters are sampled repeatedly, and used to initialise multiple optimisation runs. The contour plots indicate the objective function landscape, with the negative log-likelihood as the objective function. Note that the objective function landscape differs dependent on the hyper- parameter settings, and the model parameters are optimised in a gradient-based manner. Lower and upper bounds for a mechanistic parameter are indicated by dashed lines. Early stopping is used to determine the optimal parameter vector.

The pipeline carefully distinguishes between mechanistic parameters *θ*_*M*_, which are critical for biological interpretability, and ANN parameters *θ*_ANN_, which model components that are not well-understood or are too complex to specify explicitly. To ensure that the mechanistic parameters remain interpretable and are not overshadowed by the ANN, the pipeline supports the use of likelihood functions, constraints and priors for the mechanistic parameters as well as regularisation for the ANN. Likelihood functions, in contrast to other objective functions used in the field of machine learning, are highly relevant to allow for the computation of maximum likelihood estimates and the assessment of uncertainties.

Constraints and priors for the mechanistic parameters are used to keep them in realistic parameter ranges, push them towards plausible values and inform the sampling of initial values. Regularisation of the ANN will stabilise estimates and might prevent the ANN from inadvertently capturing vector field contributions that should be attributed to the mechanistic model. We apply weight decay regularisation to the ANN parameters, adding an L2 penalty term 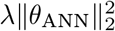 to the loss function, where *λ* controls the strength of regularisation. This regularisation discourages the ANN from becoming too complex, thus maintaining a balance between the mechanistic and data-driven components of the UDE.

The pipeline builds on parameter estimation and training methods developed for systems biology and machine learning. To account for the fact that the numerical values of parameters in biochemical reaction networks are usually vastly different, the pipeline supports the use of reparametrisation. In particular, the model supports the use of log-transformation for strictly positive parameters, thereby allowing the more efficient study of parameters spanning several orders of magnitude. For optimisers that do not support constrained optimisation, we implement a tanh-based transformation that approximates the logarithmic scale to enable bounded and scaled parameter estimation (Methods section 5, Fig. 8). Indeed, this transformation naturally enforces the requirements that mechanistic parameters are non-negative and contributes to the mitigation of issues such as vanishing gradients, thereby improving the convergence properties of the optimisation algorithm [18, 10]. The pipeline supports Maximum likelihood estimation (MLE) for identification of the maximum likelihood estimate for the mechanistic parameters and the simultaneous estimation of the noise parameters of the error model (Methods 2). Furthermore, as the parameter identification is complicated by a large parameter space and non-convex objective function, the pipeline is tailored to a multi-start optimisation strategy, that not only samples initial values for the UDE parameters *θ*_*M*_ and *θ*_*ANN*_, but also the hyperparameters such as ANN size, activation function, and optimiser learning rate. These parameters and hyperparameters are all sampled jointly to improve exploration of the (hyper-)parameter space. To further enhance the optimisation process, we incorporated common machine learning practices such as input normalisation, which improves the numerical conditioning of the problem, and early stopping, which prevents overfitting by terminating training when out-of-sample performance ceases to improve.

The pipeline leverages advanced numerical schemes to enable the study of stiff dynamical systems prevalent in systems biology [17]. Specialised solvers were required for solving the stiff systems used in this study (Supp. Table 7). In particular, we use the Tsit5 and KenCarp4 solvers within the SciML framework, ensuring efficient and accurate handling of stiff system dynamics [19, 14].

In the following sections, we utilise this pipeline to test and validate UDEs on both synthetic and real-world biological datasets. We consider the following processes, models and datasets:

– **Glycolysis model:** Glycolysis is a central metabolic pathway that describes the ATP and NADH-dependent conversion of glucose to pyruvate (see Supp. Section 10.1), and it has been extensively studied in systems biology. Our study builds upon the model by Ruoff et al. [20], which consists of seven ordinary differential equations (ODEs) exhibiting stable oscillations, depending on the parameterisation of twelve free parameters. Compared to the fully mechanistic ODE model, we replace the ATP usage and degradation with an ANN (Supp. section 10.1) that takes all seven state variables as inputs. Therefore, to recover the true solution of the data-generating process, the ANN must learn a dependency on only one of its inputs, the ATP species. We generated several synthetic data sets, allowing us to test different scenarios while providing access to the ground truth. Training (and validation) data with *t* ∈ [0, 1.5] covers a bit more than one period of the oscillation, predictive behaviour is assessed in *t* ∈ (1.5, 5]. The training data were generated using the published parameters. We considered training data with high and low (=realistic) sampling density, as well as low and high (=realistic) noise levels.
– **STAT5 dimerisation model:** STAT5A and STAT5B associate to homo- or hetero-dimers that are imported into the nucleus and modify gene expression. We study the model by Boehm et al. [21]. The model is fully specified by eight ODEs and we estimate nine mechanistic parameters. The STAT5 dimerisation example is more difficult than the glycolysis example due to the large parameter ranges spanning ten orders of magnitude, and a highly non-linear observable mapping (see Methods 4.2). We evaluate four different UDE scenarios, details can be found in the later section.

Details on the setups, including parameter bounds and reference values, are provided in the Methods section Table 3 and Supplementary Tables 4, 6.

### 2.2 Assessment of optimisation methods

Machine learning primarily aims to develop predictive models, where reproducibility of parameter estimates is often unnecessary or infeasible. Consequently, training is typically limited to a few runs, with many studies reporting results from only a single run. In contrast, systems biology studies the interpretation of the estimated parameters, motivating the pursuit of globally optimal parameter estimates. A key advantage of UDEs compared to standard machine learning approaches is the integration of prior knowledge on model structure and parameters. To exploit this advantage, optimisation methods need to provide accurate estimates for the mechanistic parameters. While multi-start local optimisation is commonly used for fully mechanistic differential equation models, this is not commonly employed for UDEs. Here we compare three parameter estimation strategies for UDEs:

– Standard Single-Start optimisation: A baseline approach starting from a single initial guess of the mechanistic parameters.
– Adapted Single-Start optimisation: An improved version incorporating enhancements such as maximum likelihood estimation (MLE), noise estimation, parameter bounds and scaling, and early stopping.
– Multi-Start optimisation: A global strategy based on the adapted single start optimisation, diversifying optimisation attempts by sampling multiple initialisations of mechanistic parameters and hyperparameters.

Using a glycolysis model with favourable data conditions (46 data points per observable, 5% noise), we evaluated the three approaches. This dataset represents a relatively simple optimisation problem, providing a suitable scenario to assess the effectiveness of the optimisation strategies. All methods employed the ADAM optimiser followed by BFGS [22, 23], as suggested in [14]. As the best model, we select the model with the best training loss (based on the dataset used for calibration) and the predictive performance was evaluated using the normalised mean absolute error (NMAE) with the true solution (test loss).

In the standard single-start optimisation approach, we initialised the mechanistic parameters *θ*_*M*_ at a single starting point and performed local optimisation. This method is commonly used in previous UDE studies, e.g., [14]. Our analysis revealed that this approach often does not lead to a successful model fit (Fig. 3a). Strikingly, even the directly observed species *N*_2_ was not fitted successfully, whereas the observed *A*_3_ species, which was approximated with a mechanistic and universal differential, closely follows the measurements.

**Fig. 3:**
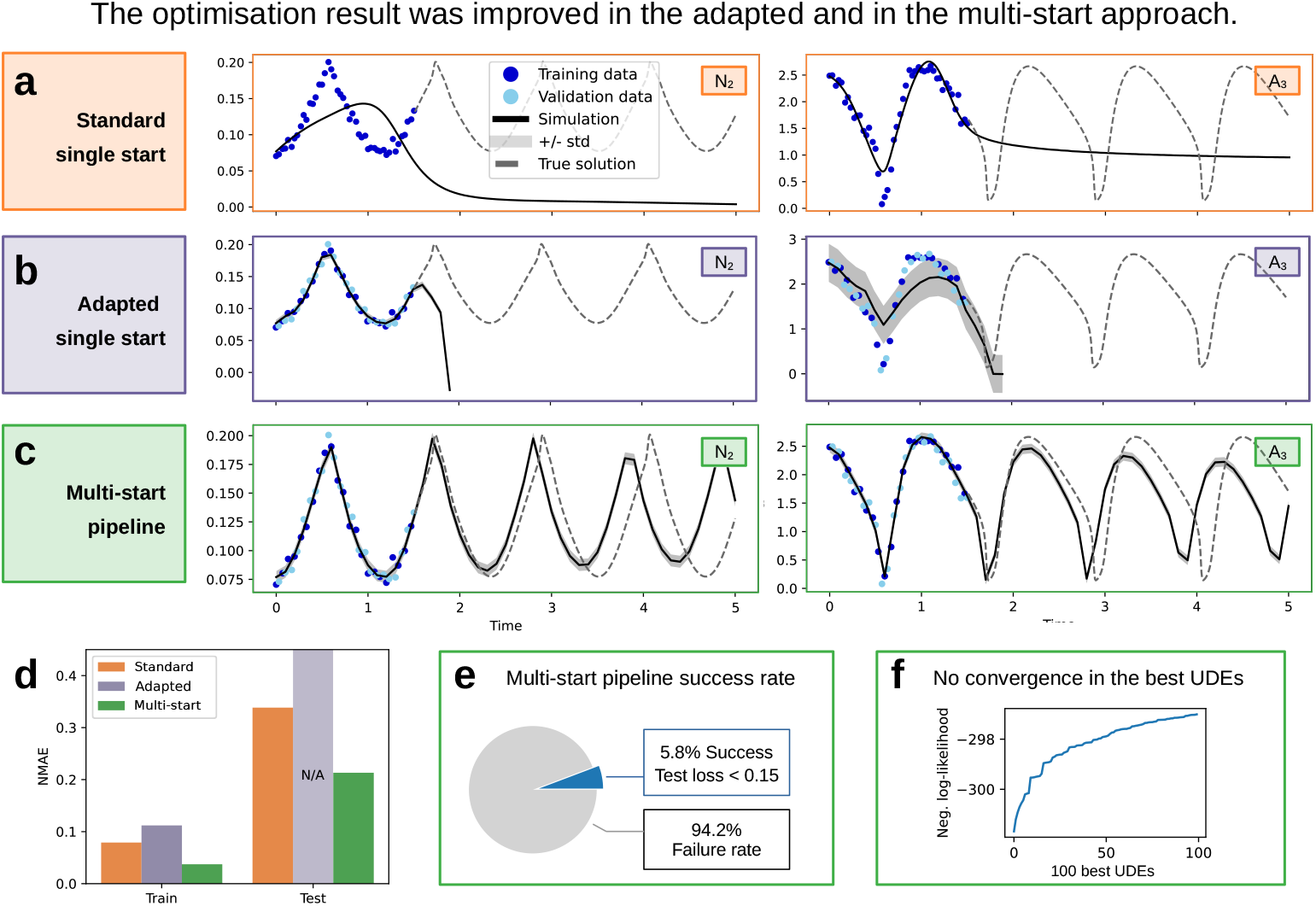
Comparison of Single- and Multi-Start Optimisation for UDE model of Glycolysis. **(a)-(c)** (left) Best fit on training data and corresponding prediction achieved by standard single-, adapted single-, and multi-start approaches for the data set with 46 data points and 5% noise. **(d)** Comparison of the NMAE on the training and test set by approach. The test loss for the adapted approach is not computable due to the simulation failure (b). **(e)** The success rate (Test loss < 0.15) for UDEs in the multi-start pipeline. **(f)** Waterfall plot of the training loss for the 100 best optimisation runs of the multi-start approach.

In the adapted single-start approach leverages the MLE, transformed parameters, as well as a train-validation data split. Despite these improvements, the adapted single-start optimisation did not achieve a satisfactory model fit (Fig. 3b). Due to the noise modelling in the adapted approach, we can still retrieve informative confidence levels for the simulations (Fig. 3b, right). Simulation fails in the testing regime due to a blow-up in the *N*_2_ species. This suggests that simply refining the single-start method is insufficient for successful estimation of UDE parameters in this context.

The multi-start optimisation strategy involves initiating multiple optimisation runs, realised as initiating model optimisation 10 000 times from diverse hyperparameter settings and starting points for *θ*_*M*_ . This approach significantly improved the optimisation results, leading to a successful model fit (Fig. 3c). Comparing the three approaches quantitatively (Fig. 3d), the multi-start optimisation achieved better training and test errors compared to both single-start methods. Therefore, the multi-start approach improves the fit to the measurements, and also enhances predictions.

Based on the distribution of test losses and on visual validation, we set a threshold of 0.15 for the test error to distinguish a successful fit. Examples of a successful and failed model fits according to the criterion are shown in Supplementary Fig. 10f-h. Only a fraction of the multi-start runs resulted in a fit with a good test metric (Fig. 3e) and we did not find convergence in the training objective or mechanistic parameters (Fig. 3f, Fig. 9). The lack of convergence demonstrates the challenges in the optimisation landscape of UDEs, even under favourable data conditions.

Despite the increase in computation time for a higher number of starts, our results emphasise the necessity of a global optimisation strategy, such as multi-start optimisation, for inferring a useful process description and the UDE parameters. The standard single-start approach is insufficient, and additional enhancements do not guarantee success. The inherent complexity of the UDE optimisation landscape necessitates extensive exploration of the parameter space to identify optimal solutions.

### 2.3 Impact of data density and measurement noise

Given the critical role of the optimisation procedure, we evaluated the impact of data properties on the performance of UDEs. Specifically, we focused on data sparsity and noise levels, common challenges in systems biology due to technical limitations. We examined twenty data settings, varying in density (8, 16, 31, 46, and 61 data points per observable) and noise levels (5%, 10%, 20%, and 35%). Measurements cover a single oscillation of the glycolysis model. For each data setting, we applied the previously described multi-start optimisation strategy with 10 000 starts.

For all data settings, the UDEs with the lowest training loss achieved a good description of the training data, and the estimated noise was generally informative, deviating at most 50% from the true value (Fig. 4e, Supp. Fig. 12). Our analysis revealed that the data information content, in terms of sparsity and noise, was decisive for the inference of accurate process descriptions.

**Fig. 4:**
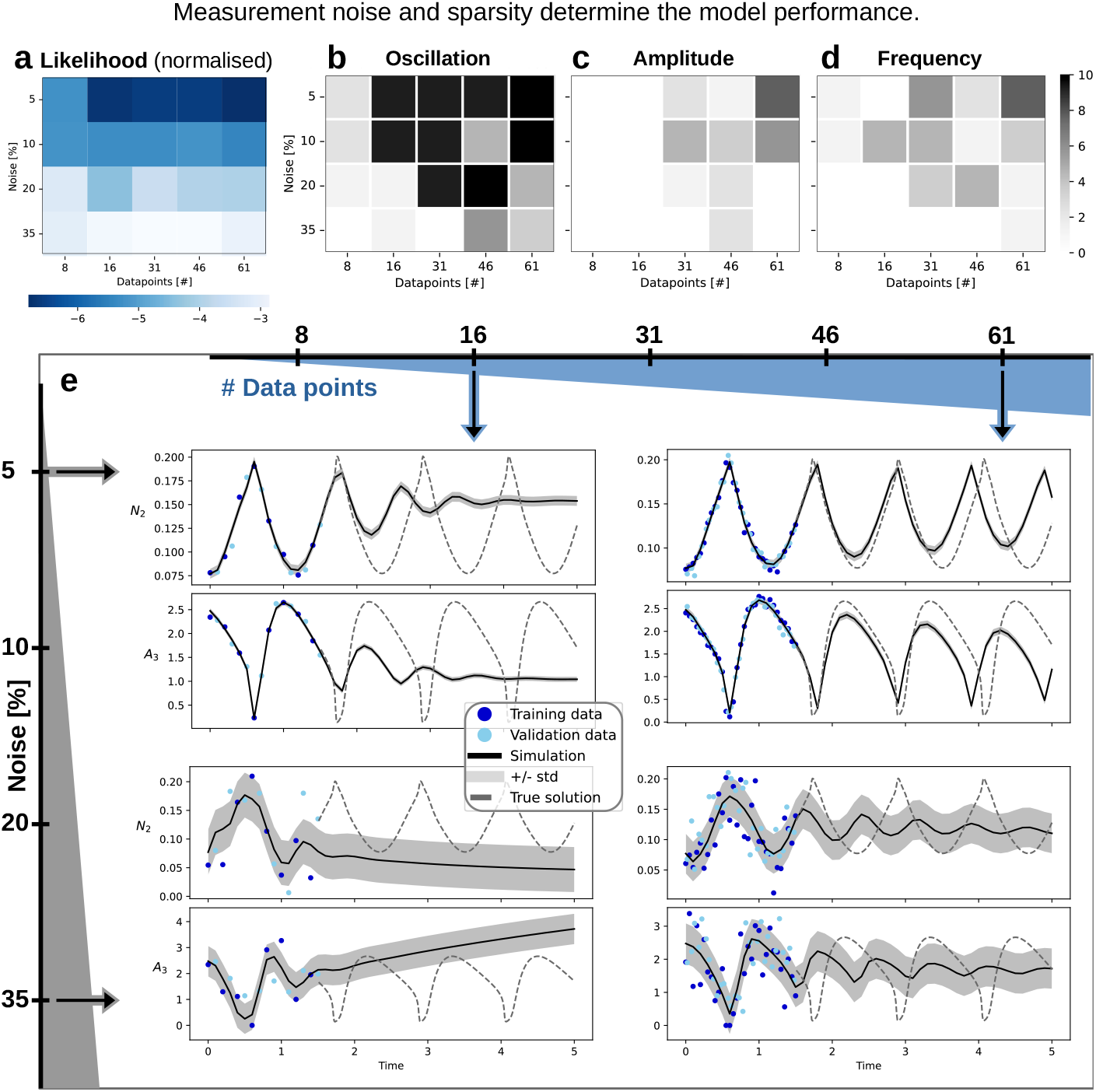
Assessment of the impact of data density and noise levels on the performance of UDE model of Glycolysis. **(a)** Training loss for all data settings: The colour indicates the negative log-likelihood of the best UDE model by noise/sparsity setting, normalised by the number of data points used for training. **(b)-(d)** The number of fits in the 10 best fits (by training loss) per noise/sparsity setting that recover (in predictions) **(b)** oscillatory behaviour, **(c)** amplitude, or **(d)** frequency close to the true solution. **(e)** Best fits (= best training loss) from four different noise/sparsity settings: 16 (left column) and 61 (right column) data points per observable; and low noise (upper row, 5%) or high noise (lower row, 35%).

Higher noise levels significantly degraded the quality of fit and prediction performance (Fig. 4a), while data sparsity had a particularly pronounced impact in the low-data regime (e.g., 8 vs. 16 data points). A good quantitative predictive performance was restricted to the easier data settings with higher data density and lower noise (Fig. 4e, Supp. Fig. 12). To assess whether UDEs allow for the reconstruction of qualitative model properties from limited data, we analysed the ability of the trained model to recover sustained oscillations and match the amplitude and frequency of the true solution (Fig. 4b-d). Under poor data conditions, UDEs often recovered dampened rather than sustained oscillations. Oscillations with quantitatively informative predictions were observed only in the best-case scenarios (e.g., 5% noise and 61 data points per observable, Fig. 4e, top right; or 46 data points, Fig. 3c), at least in the initial unobserved cycles.

Interestingly, we did not observe a consistent improvement of our results with decreasing noise and increasing data points (Fig. 4a-d), underscoring that convergence is not assured, and the optimisation landscape remains challenging under varying data conditions. Notably, models with better predictions were present across all data settings when evaluated by test loss, with the best overall test loss consistently improving as noise decreased and data density increased (Supp. Fig. 11).

### 2.4 Effect of hyperparameters and regularisation

As we observed that training loss is not a reliable indicator of successful model fit and prediction, we investigated whether hyperparameter settings, particularly regularisation, affected the results and whether certain settings could guarantee improved predictions.

Our re-analysis of the results generated for the previous evaluation showed that successful fits were achieved across a wide range of hyperparameter settings (Fig. 5a), indicating a certain degree of robustness. Most hyperparameter settings were approximately equally likely to lead to a successful fit. There were slight tendencies against the use of input normalisation and towards higher initial learning rates in the ADAM optimiser among successful runs. There was no clear tendency in the choice of activation function and ANN size, suggesting that, as long as the ANN is flexible enough to approximate the missing dynamics, modelling can be successful. Increasing the ANN dimension led to a slight increase in optimisation time (Supp. Fig. 14), indicating a disadvantage rather than a benefit to using a high-capacity ANN. Similarly, the successful models were initialised with mechanistic parameters from the entire parameter ranges, with low tendency for specific sampled values to enhance the chance for success (Fig. 5b, Supp. Fig. 13). The (hyper-)parameter vectors that serve as suitable initialisations for optimisation are arranged on a problem-specific manifold that is difficult to ascertain *a priori*. Taken together, the results suggest that the multi-start strategy is able to sample from this manifold, unlike the standard or adapted strategies, and is relatively robust to problem-specific features like oscillating data.

**Fig. 5:**
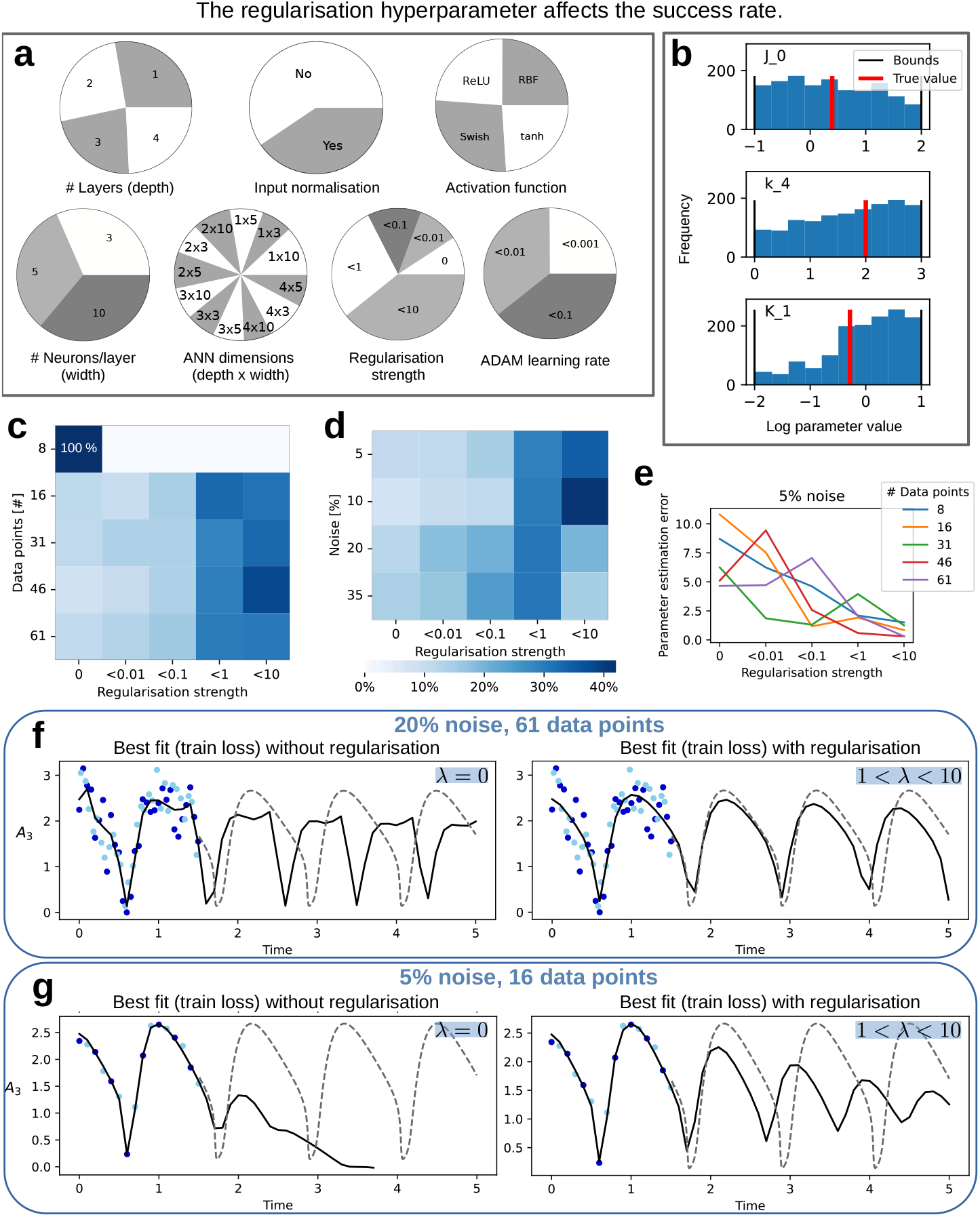
Impact of hyperparameters and regularisation on modelling success. **(a)** Hyperparameters of all successfully fitted models (test loss *<* 0.15). Each pie represents one hyperparameter with the pie’s fractions corresponding to the number of successful fits with a specific setting. The continuously sampled hyperparameters (learning rate, regularisation strength) were binned, and the labels show the upper bound. **(b)** Initial values for three of the mechanistic parameters, for all successfully fitted models (test loss *<* 0.15). **(c, d)** Percentage of successful fits (test loss *<* 0.15) by regularisation strength, as percentage of the overall successful fits per **(c)** dataset size, and **(d)** noise level, i.e., each row sums to 100%. **(e)** Parameter estimation error for the best model (lowest training loss) by regularisation strength, shown for all data sets with 5% noise. **(f, g)** Best model fits (lowest training loss) for different data sparsity and noise levels, and regularisation strength.

Importantly, the regularisation strength (weight decay parameter *λ*) had the strongest effect on optimisation success. The number of successful fits increased with higher regularisation strength, with successful fits being more than four times as likely with 1 *< λ <* 10 compared to no regularisation. The evaluation of the impact of regularisation across different data densities and noise levels (Fig. 5c-d) revealed that the majority of successful UDEs were trained with *λ >* 0.1, except in the lowest data availability setting (8 data points per observable), where only one successful fit was obtained, which was without regularisation. Similarly, for different noise levels, the regularisation settings with the highest fraction of successful fits were 1 *< λ <* 10 for noise levels 5% and 10%, and 0.1 *< λ <* 1 for noise levels 20% and 35%. To assess the regularisation effect on recovering the mechanistic parameters, we compared the sum of squared errors between the estimated and true parameters in log space. We found that regularisation can substantially improve the accuracy of the inferred parameters, most pronounced in the easier data settings (Fig. 5e, Supp. Fig. 17).

The positive effect of regularisation was evident across data set sizes and noise levels. In easier data settings, appropriate regularisation led to quantitative improvements in prediction quality (Fig. 5f). In more challenging settings, regularisation mitigated issues such as exploding dynamics, favouring the recovery of weak oscillations (Fig. 5g; see also Supp. Fig. 15, 16). Accordingly, well-tuned regularisation not only increased the number of successful fits but also improved the quality of models based on training error. To assess whether the ANN approximates the true mechanism, we used an evolutionary algorithm for symbolic regression [24]. However, even with regularisation, the learned ANN mechanisms did not appear similar to the true mechanism (Supp. Table 5). This is potentially due to non-identifiability as the input species to the ANN share similar, oscillatory dynamics.

In summary, our results demonstrate that data density and noise levels critically influence the optimisation success of UDEs in systems biology applications. The use of multi-start optimisation is necessary but not sufficient; appropriate regularisation plays a key role in balancing the mechanistic and ANN contributions, leading to improved extrapolation, predictive performance and parameter estimation. However, the ANN component may remain non-identifiable despite these improvements.

### 2.5 Real-world application: STAT5 dimerisation

To corroborate the findings from the glycolysis application in a real-world context, we applied the UDE approach to a parameter estimation problem for the STAT5 dimerisation process, a key mechanism in cellular signal transduction [21]. We assessed the performance of the UDE approach across four scenarios, each simulating the absence of specific mechanisms from the original model. In these cases, an ANN approximated the missing mechanisms:

– *Scenario 1 - pApA export kinetics:* We investigate a UDE with one unknown mechanism, here the export and dissociation of the nuclear pApA dimer (`nucpApA`).
– *Scenario 2 - pApB differential:* The right-hand side of the `pApB` species is fully approximated by an ANN.
– *Scenario 3 - Augmented export reactions:* In *Scenario 3* we use the full mechanistic model and introduce additional species with fully universal dynamics, `K`_*A*_, `K`_*B*_, `K`_*AB*_, to augment the dynamics describing the export reactions (Fig. 6g).
– *Scenario 4 - Observable STAT5A/B ratio*: The ANN replaces the observable mapping between the measurements and dynamic system.

**Fig. 6:**
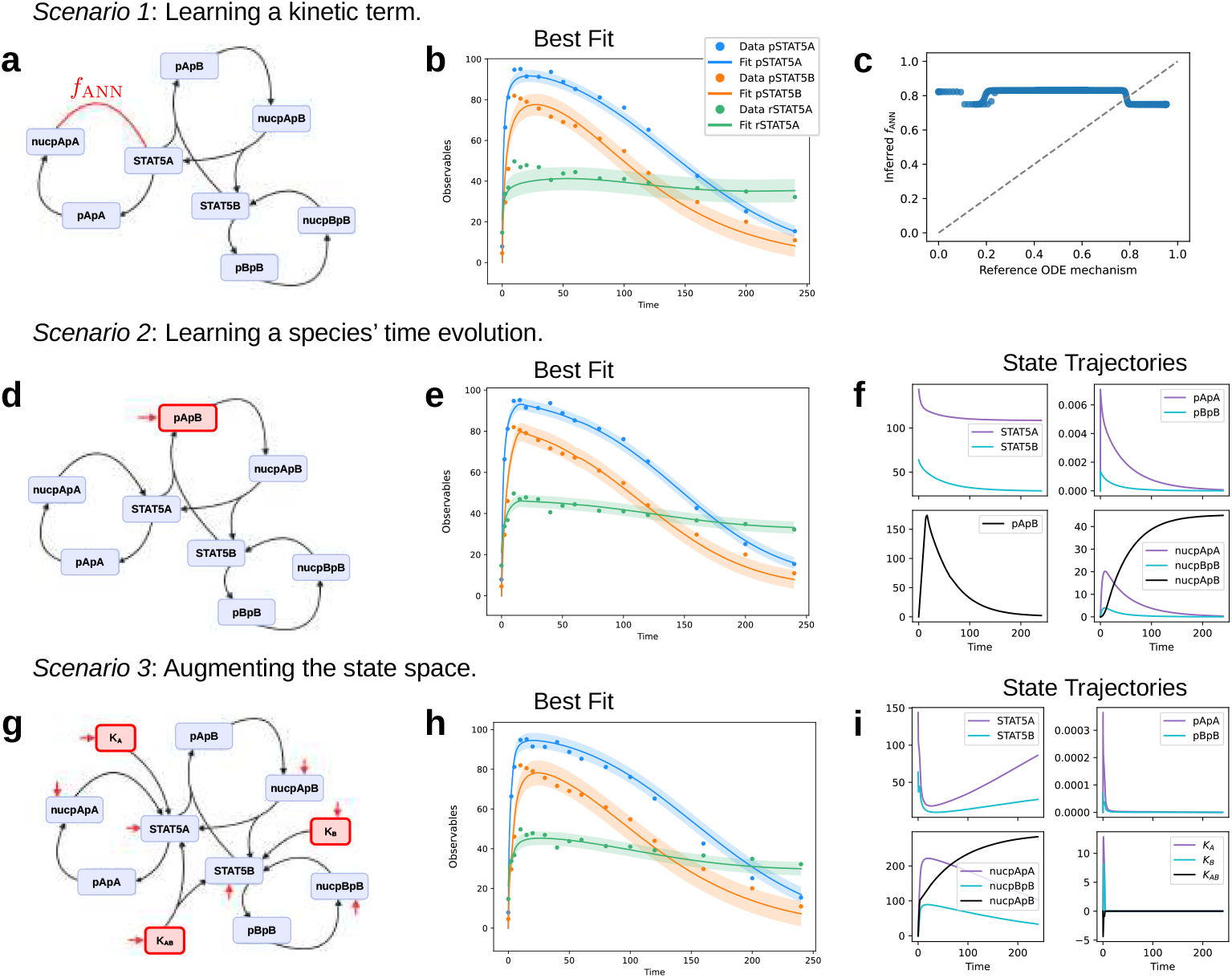
Evaluation of UDEs in different settings. **(a, d, g)** Model schematics for the UDEs used in each *Scenario*. Red elements indicate ANN source terms. **(b, e, h)** Best UDE fit for each *Scenario*. Error bands are fit *±* 1 std. dev. (estimated). **(c)** Comparison of the inferred *f*_*ANN*_ and corresponding term in the reference ODE, on the same input. A variety of inputs were taken from the reference ODE and optimised UDE simulations. **(f, i)** Simulation trajectories for the unobserved species, from the best UDE in *Scenarios 2* and *3*.

We used a similar multi-start training procedure as for the glycolysis model, refer to Supp. section 10.2 for the details. In contrast to the Glycolysis example, this analysis relied on actual measurement data, meaning that no independent test data are available. The measurements were split into training and validation set at a 4:1 ratio to enable early stopping. Based on the findings for the Glycolysis model, we fixed the dimension and activation function of the ANN in *Scenarios 1-3* and use only the regularisation strength, the ADAM learning rate initialisation and the input normalisation as tunable hyperparameters (Methods section Table 3).

### 2.6 A biological constraint increases interpretability

The pApA export reaction in *Scenario 1* is approximated by an ANN that takes the full state vector as input and modifies the dynamics of `nucpApA` and `STAT5A` (Fig. 6a). To ensure mass conservation, we imposed a constraint by transforming the ANN output such that the sum of the outputs equals zero. This constraint operates at the level of the stoichiometric matrix, preserving mass balance for fluxes between `nucpApA` and `STAT5A`.

The multi-start optimisation strategy yielded a reasonable fit for the complex STAT5 dimerisation problem under this mass conservation constraint (Fig. 6b). Although the peak was less pronounced than in the reference model, especially for the observable `rSTAT5A`, the estimated noise appropriately captured the uncertainty. Among the 3 000 optimised UDEs for *Scenario 1*, only few achieved a reasonable fit by visual assessment. While each hyperparameter setting was shown to lead to a good fit, and a high learning rate was more likely to lead to a good fit (Supp. Fig. 18). Regularisation appears to have limited importance in this scenario, likely because ANN’s influence was constrained to modifying only two species and was consistent with mass conservation laws.

Despite good agreement between model simulations and measurements, the estimated mechanistic parameters *θ*_*M*_ did not match the published values for the STAT5 dimerisation model ([21], Supp. Table 6). Instead, the UDE models found different local optima for this parameter estimation problem, potentially due to parameter non-identifiabilities introduced by the ANN and the high complexity of the optimisation landscape.

Symbolic regression of the inferred ANN yielded a cautiously interpretable result. The recovered functional form, 3.96 ·10^−4^ · *x*_6_ + 0.8 deviated from the true functional form, 6.17 · 10^−2^ · *x*_6_, only in a constant term. The `nucpApA` species (*x*_6_) was correctly identified as the sole relevant input among all species of the over-parametrised model (117 free parameters). Fig. 6c shows the respective contributions of the reference ODE term and the inferred ANN, where the constant term dominates in the UDE case.

Overall, the positive fitting result for *Scenario 1* of the realistic STAT5 dimerisation model can be attributed to the constrained flexibility of the ANN, which reduces the relevance of the naïve parameter regularisation for this setting.

### 2.7 ANNs as the state derivative replace and augment the ODEs

In *Scenario 2*, the right-hand side of the `pApB` species was fully approximated by an ANN, which used all species and the time-varying concentration of the erythropoietin stimulus (`BaF3_Epo`) as inputs (Fig. 6d). Parameter inference resulted in a good fit to the measurements (Fig. 6e), with a marked improvement in the objective function value compared to ODE optimisation (negative log-likelihood 81.1 vs. 138.22), using *λ* = 0.0145. Simulations showed that the `pApB` species exhibited a time evolution that was qualitatively similar to the homodimers `pApA` and `pBpB` (see Fig. 6f). The `pApB` concentration peaked slightly later but reached a distinctly higher value, consistent with the behaviour of the ODE model, where `pApB` concentrations are orders of magnitude higher than those for `pApA` and `pBpB`. These results demonstrate that a UDE with a flexible ANN fully describing the derivative of a non-observed species can effectively infer dynamic behaviour.

The improvement in the likelihood could indicate that the mechanistic terms are suboptimal for describing the dynamics of the hetero-dimer `pApB`. However, the lack of biologically informed constraints in the UDE model limits its validity, as it does not guarantee biologically sound behaviour. Interestingly, regularisation provided no clear advantage in this scenario (Supp. Fig. 18). As the input species to the ANN move on different orders of magnitude, we evaluated if the result can be improved further by input normalisation (Supp. Fig. 18). Interestingly, we found a slight tendency against input normalisation, likely because differences in the magnitudes of input species carried informative signals. Normalising these inputs may have inadvertently removed critical information, leading to a negative effect on model performance.

Building on the hypothesis that UDEs can identify areas of incomplete mechanistic knowledge, we used UDEs to explore relevant, missing components for an existing mechanistic model in *Scenario 3*. In the original STAT5 dimerisation model, dimer export from nucleus, dimer dissociation and dephosphorylation are simplified into a single mass-action term. This simplification might limit the accuracy, so we investigated additional mechanisms by introducing the augmentation variables *K*_A_, *K*_B_ and *K*_AB_, modelled using an ANN. The augmented UDE incorporates unknown, potentially relevant species, thus extending the published mechanistic model. In this scenario, we build on a consistent ODE model without missing interaction terms. Therefore, as an alternative form of balancing mechanistic and ANN contributions, we applied a two-stage optimisation strategy. First, the mechanistic model was optimised independently. Then, the full UDE model was optimised starting from the values estimated for the mechanistic parameters.

Our augmented system in *Scenario 3* achieved a fit that captured the overall dynamics of the measurements (Fig. 6h). The best-performing model utilised regularisation (*λ* = 0.21), leading to augmentation species that exhibited pulse-like behaviour (Fig. 6i). This pattern mirrors the transient activity spikes observed in the `pApA`, `pApB` and `pBpB` species. However, the simulated trajectories for the augmentation species revealed biologically implausible behaviour, with *K*_AB_ concentrations becoming negative. This issue could be addressed by introducing tailored constraints to enforce non-negativity [15], although such constraints might impede the implicit flexibility of the ANN. These findings highlight a critical trade-off in UDE design: flexible ANNs are valuable for uncovering novel mechanisms, but constraints are often necessary to ensure adherence to biological principles.

Despite a marked improvement in likelihood, the AIC and BIC model selection criteria indicate that the UDE model does not improve on the ODE model due to the high number of estimated parameters (Tab. 1).

**Table 1:** Akaike information criterion (AIC) and Bayesian information criterion (BIC) calculated based on the negative log-likelihood (NLL) of the reference ODE model and the best UDE model.

| Method | NLL | # Parameters | # Data points | AIC | BIC |
| --- | --- | --- | --- | --- | --- |
| ODE | 138.22 | 9 | 48 | 294.44 | 311.28 |
| UDE | 86.88 | 153 | 48 | 479.76 | 766.0 |

### 2.8 UDEs incorporate additional measurements

Finally, we propose to use an ANN for the observable mapping between the measurements and dynamic system, bypassing the numerical integration (see Fig. 1). The best fit achieved by a UDE without regularisation exhibited clear signs of overfitting (Fig. 7b), which was mitigated when applying regularisation (Fig. 7c). Comparing the outputs of the inferred ANN to the mechanistic observable mapping, on the same inputs, revealed notable differences (Fig. 7f).

**Fig. 7:**
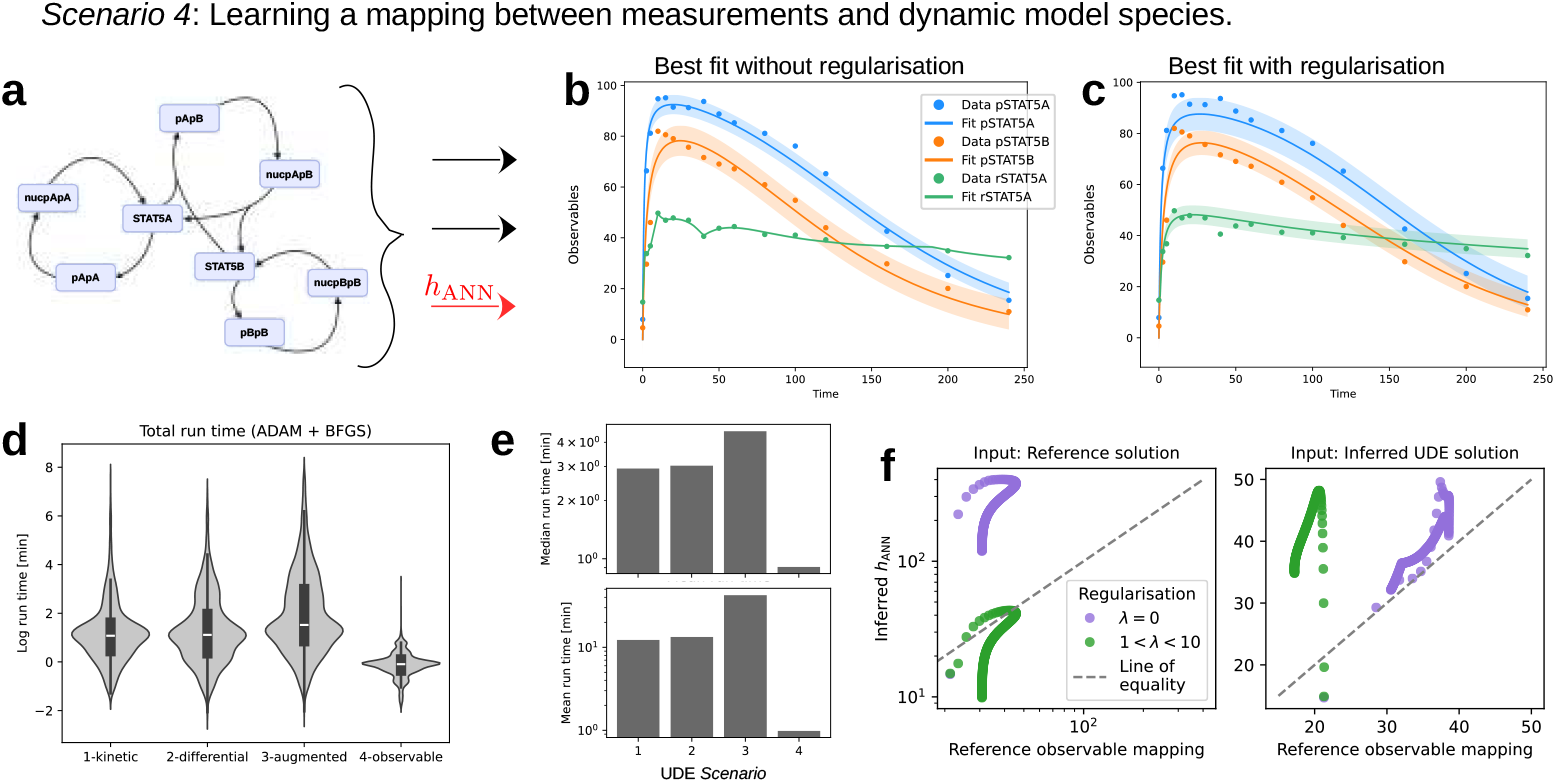
Evaluation of UDEs with approximated observable mapping. **(a)** Schematic of the UDE setup for *Scenario 4* where black elements correspond to mechanistic ODE terms and red elements indicate ANNs. **(b, c)** Best fitted UDE for *Scenario 4* without and with regularisation. **(d)** Total optimisation time for each trained UDE per *Scenario*. **(e)** Mean and median total optimisation time by *Scenario*. **(f)** Comparison of the inferred *h*_*ANN*_ and corresponding observation function in the reference model, on the same input. A variety of inputs were taken from the reference ODE (left) and optimised UDE (right) simulations.

When using the reference solution as inputs, the regularised UDE aligned better with the magnitude of the original observable mapping (Fig. 7f, left). However, on the UDE-inferred solution, the non-regularised ANN exhibited better agreement with the mechanistic observable mapping (Fig. 7f, right). Despite the positive effect of regularisation on the best fit, there was no consistent improvement in the approximation of mechanistic observable mapping.

The ANN’s placement significantly influenced runtime. When applied to the observable mapping rather than the differential equation, runtime decreased substantially. Optimisation for *Scenario 4* averaged less than one minute per UDE, compared to approximately 12 and 45 minutes for *Scenarios 1-3*. This reduction may be due to the simpler evaluation of parameter gradients in the observable mapping, unlike ANN parameters embedded in the dynamic equations, which require differentiable programming such as adjoint sensitivity analysis. However, the difference in median computation times was smaller (a factor of 0.2–0.3), suggesting a strong impact of outliers corresponding to optimisation runs with very high computation times (Fig. 7e).

To corroborate our earlier findings, we analysed the hyperparameters of the UDEs. In the absence of a test data set, we used the *χ*^2^ test to identify UDEs in the 95% confidence region around the best fitted UDE for evaluating the hyperparameters leading to a good fit. Similar to the Glycolysis UDEs, we found no reliable trends favouring specific hyperparameter settings (Supp. Fig. 18).

## 3 Discussion

The integration of Universal Differential Equations (UDEs) into semi-mechanistic modelling offers a transformative approach to addressing complex biological systems. By combining the flexibility of neural networks with the interpretability of mechanistic models, UDEs bridge the gap between data availability and mechanistic knowledge. Their success, however, hinges on efficient optimisation approaches that yield accurate and interpretable results. Our study evaluates optimisation strategies, as well as the sensitivity of UDE performance to dataset size and noise. We observed that multi-start optimisation, which is commonly used for parameter inference in systems biology, is also required for UDEs. The objective function landscape appears to be usually multi-modal, and single start methods were not able to escape the modes but showed a dependence on the starting point. The number of required optimisation starts tends to decrease with lower noise levels and higher data availability. Leveraging parameter estimation techniques from systems biology, we successfully applied UDEs in realistic systems biology settings characterised by sparse data and substantial measurement noise.

The biologically informed scaling approach, implemented as a tanh-based transformation, facilitated the exhaustive exploration of parameter space and estimation of biologically meaningful parameters. Efficient optimisation was central to successful UDE modelling. Employing efficient stiff solvers rendered a computationally-intensive multi-start optimisation strategy feasible. The maximum likelihood approach effectively accounts for the significant measurement noise, guiding a statistically motivated, robust model selection over merely achieving a close fit to noisy data. By incorporating realistic noise models, this method eliminates the need for output scaling or additional weighting in the objective function, thus streamlining the optimisation process. The challenge imposed by a complicated observable mapping and limited data is especially apparent in the real data application example, where only few models resulted in a sensible fit and achieving convergence remained challenging. Our research highlighted the challenge of parameter non-identifiability and the resulting pronounced uncertainties in UDEs, which can complicate the biological interpretability of results [25]. This emphasises the need for a comprehensive exploration of the parameter space through a global multi-start optimisation strategy.

The hyperparameter tuning was facilitated through the multi-start approach and improved diversity of the optimisation start points. As shown for the glycolysis application, our results indicate no consistent dependency on a particular setting for ANN width, depth or activation function. Our study emphasises the importance of pragmatic hyperparameter selection, driven by considerations such as computational efficiency and mitigating the risk of overfitting. Notably, our findings challenge prevailing assumptions in machine learning-based modelling by showing no evidence for the necessity of input normalisation when working with variables that vary by orders of magnitude.

Overfitting remains a persistent challenge in UDE modelling, particularly in the absence of a test set to validate generalisation. The over-parameterised ANN component consistently hindered parameter identifiability, thus complicating the modelling process. To address these challenges, regularisation techniques and biologically-informed constraints proved essential. Regularisation through the generic weight decay method enabled smooth, realistic fits to the measurements of the STAT5 dimerisation process and proved beneficial for enhancing predictive performance in the glycolysis application. However, it was insufficient for reliably recovering the mechanistic parameters of the known network components. This highlights the need for advancing regularisation methods that balance the flexibility of neural network components with the interpretability of mechanistic terms. As demonstrated for the approximation of the `pApA` export in the STAT5 dimerisation model, a biologically-informed mass conservation constraint facilitated the recovery of realistic fits. Future research should prioritise refining regularisation techniques to improve generalisation, preventing overfitting, and strike a balance between the contributions of the UDE components. The knowledge embedded into the mechanistic terms should be preserved, while the data-driven inference by the universal approximator should remain maximally flexible. Regularisation and biologically consistent constraints will be critical for ensuring biologically meaningful behaviour in UDE models, as also supported by previous studies, such as [15].

Our study highlights several challenges of using UDEs in systems biology. Yet, it also pinpoints the tremendous potential. The analysis of the STAT5 dimerisation model revealed that a reasonable UDE fit was possible for different scenarios. ANNs were successfully used to model reactions corresponding to kinetic rates, and dynamic variables, demonstrating their utility in systems modelled with stoichiometric matrices or differential equations. Another prominent example is using UDEs to model inputs to the system, or time-dependent parameters [25], further showcasing their adaptability in augmenting existing systems of ODEs. Moreover, ANNs hold promise as observable mappings, linking experimental data with dynamic model variables. Machine learning approaches, with their proven capacity to handle large datasets, offer a pathway for leveraging additional data sources for dynamic modelling. This capability can be extended to integrate high-dimensional omics datasets, paving the way for richer insights into biological systems. The potential of UDEs with ANNs in the observable mapping could be further enhanced through hierarchical optimisation methods, which have a significant impact on model performance [26].

In conclusion, UDEs represent a promising tool for tackling challenging modelling problems, offering a pathway to novel insights into complex biological systems. Our evaluation pinpoints the challenges during parameter estimation and highlights the central role of (hyper)-parameter space exploration and regularisation in overcoming them. With continued methodological development, particularly in regularisation and optimisation, UDEs can provide interpretable and biologically meaningful predictions that bridge the divide between data-driven and mechanistic approaches.

## 4 Methods

### 4.1 Universal differential equations

In this study, the dynamics of *n*_*s*_ biological species 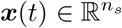 (state variables) are described through a system of coupled ODEs. The biological entities are partially or indirectly observed, thus necessitating an observable model to map the ODE state space to *n*_*o*_ variables 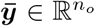 that can be measured. Universal differential equations (UDEs) combine prior knowledge about the mechanistic behaviour of a system with universal function approximators, such as artificial neural networks, to describe the dynamics of unknown mechanisms [27]. We consider the scenario where the output of a neural network is added as an additional term (summand). Hence, we define a UDE as

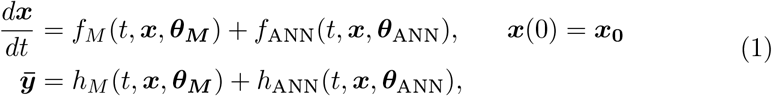

where *f*_*M*_, *h*_*M*_ and *f*_ANN_, *h*_ANN_ are, respectively, the mechanistic and neural network components of the equation system with mechanistic parameters *θ*_*M*_ and neural network parameters 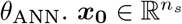 is the initial condition.

### 4.2 Problem scenarios

The UDEs used are based on two previously published biological models: The glycolysis model [20] and the STAT5 dimerisation model [21]. In the following, we define these two models along with their UDE scenarios, specify the noise model and the data.

#### Glycolysis model

The differential equations of the original Glycolysis model are:

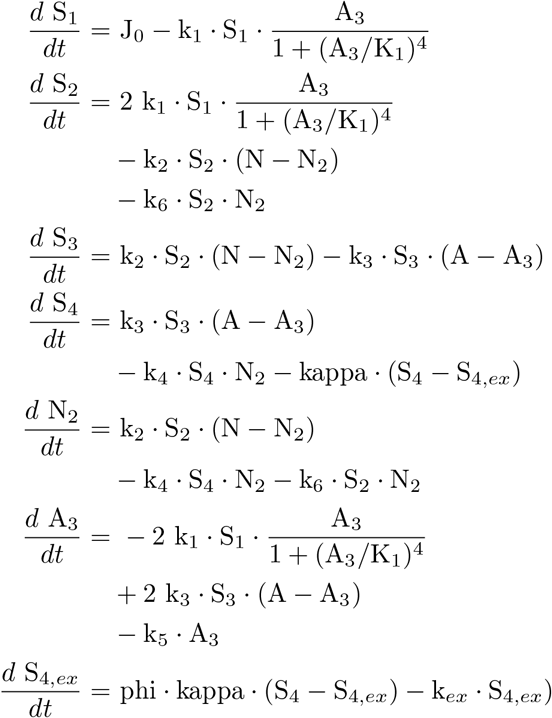

with directly observed species N_2_ and A_3_.

The ODE model was implemented in the Systems Biology Markup Language (SBML, [28]). Data generation was facilitated through simulation with the Advanced Multi-language Interface for CVODES and IDAS (AMICI) [29]. The full ODE model was optimised for reference; the parameter estimation problem was specified in the PEtab format [30] and performed using the Python toolbox pyPESTO [31]. The estimated mechanistic parameters, their bounds and the scale used for optimisation are listed in Table 4.

#### Glycolysis UDE setting

We assume that the ATP usage is unknown, i.e. remove the term −k_5_ *·* A_3_ in the right-hand sight of 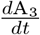. We approximate this process with a neural network that modifies the *A*_3_ dynamics (i.e. one output neuron) and provide all seven state variables as inputs.

#### STAT5 dimerisation model

The parameter estimation problem was taken from the PEtab benchmark collection [32]. The differential equations of the original STAT5 dimerisation model are:

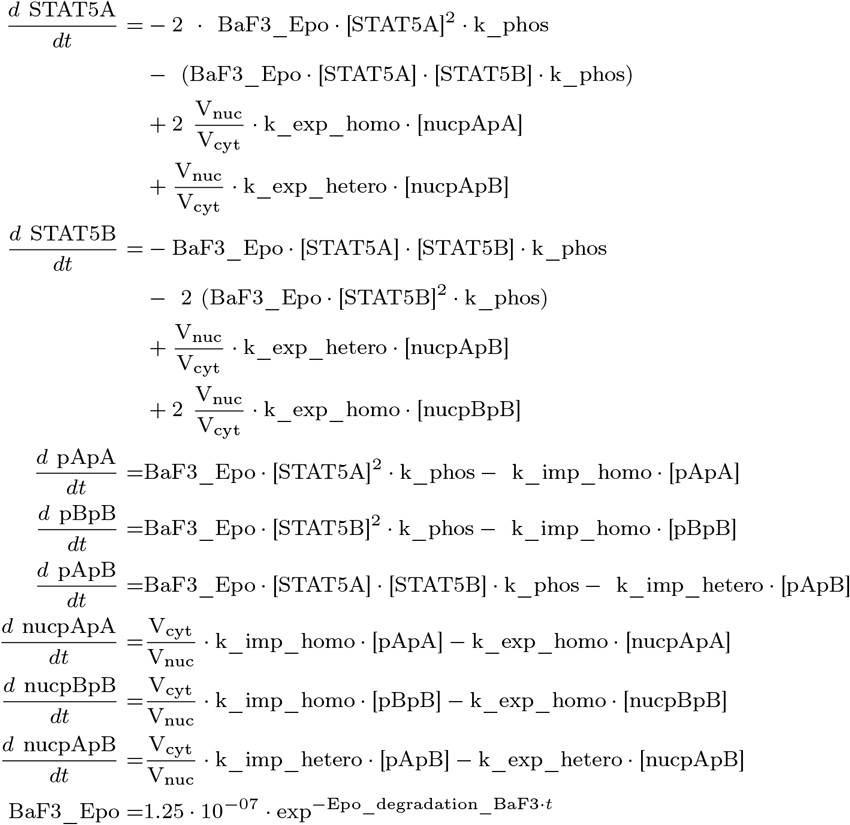

where V_cyt_ = 1.4 and V_nuc_ = 0.45. The observable mapping is given by:

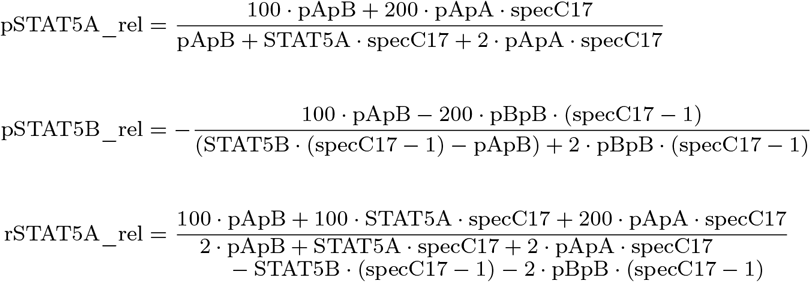

where specC17 = 0.107.

The full ODE model was optimised for reference using the AMICI simulator, the PEtab format and the pyPESTO toolbox [29, 30, 31]. The estimated mechanistic parameters from the original publication [21], their bounds and the scale used for optimisation is listed in Table 6.

#### STAT5 dimerisation UDE

**Scenario 1** In this scenario, the conversion reaction between `nucpApA` and `STAT5A` was replaced by an ANN. We assumed mass conservation by keeping the factor 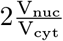 in the right-hand side of `STAT5A` and using opposite signs for the ANN in the dynamic equations of `nucpApA` and `STAT5A`, yielding

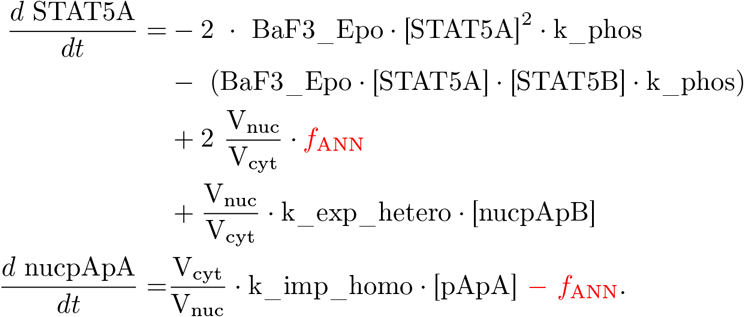

The dynamic equations of the other states and the observable mapping remain as defined for the Glycolysis model. We provide all state variables as inputs to the neural network.

#### STAT5 dimerisation UDE

**Scenario 2** In this scenario, the rate of change of the `pApB` species is entirely learned by an ANN, yielding a new dynamic equation for `pApB`:

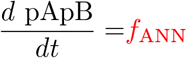

We provide all state variables and the time-dependent value of BaF3_Epo as inputs to the neural network.

#### STAT5 dimerisation UDE

**Scenario 3** An ANN describes the time evolution of the newly introduced augmenting species, and the augmenting species are converted into the monomer species `STAT5A` and `STAT5B` in the new mass action terms (bold). Instead of replacing ODE terms, the differential equations are extended by new terms. The updated differential equations read as follows, where “…” stands for the original terms of the respective differential equation:

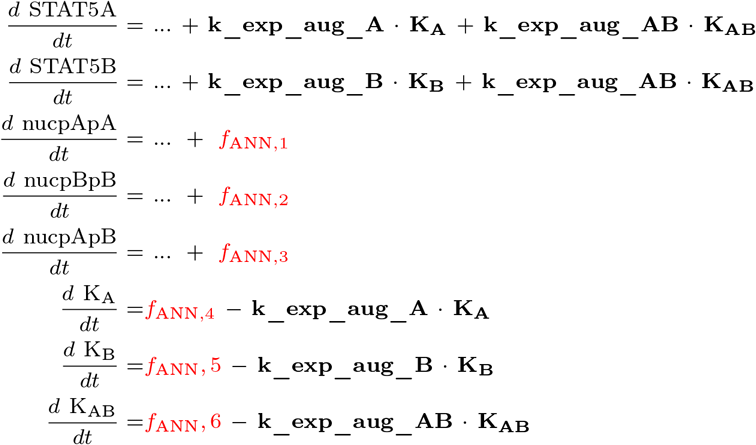

We provide the state variables `nucpApA`, `nucpApB`, `nucpBpB` and the three augmenting species as inputs to the neural network. The neural network has 6 output neurons.

#### STAT5 dimerisation UDE

**Scenario 4** In this scenario, we approximate the observed relative abundance of STAT5A by an ANN:

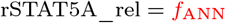

Hence, the ANN is not part of the dynamical system. We provide all state variables as inputs to the neural network.

#### Noise model

For all problems, we assume additive normally-distributed measurement noise, i.e.

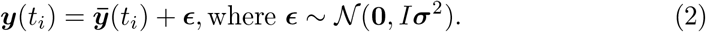

where 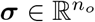 describes the observable-specific standard deviation. We estimate ***σ*** alongside *θ*_ANN_ and *θ*_*M*_ and assume ***x***_0_ to be known.

#### Data

For the glycolysis UDE, the data is a noise-corrupted realisation of the simulation of the observables defined in Section 4.2. We investigated 4 noise and 5 sparsity settings, as defined in Table 2. For the models describing STAT5 dimerisation, we use the real-world measurements presented in [21]. The train-validation split is implemented according to Table 2.

**Table 2:** Overview of train-validation split per problem scenario. The known initial condition for the Glycolysis problem was not considered for the splitting, but is counted in the total number of data points.

| Problem | Noise [%] | Data points [#] | Train-validation split |
| --- | --- | --- | --- |
| Glycolysis | [5, 10, 20, 35] | 8 | 4:3 |
| Glycolysis | [5, 10, 20, 35] | 16 | 7:8 |
| Glycolysis | [5, 10, 20, 35] | [31, 46, 61] | 1:1 |
| STAT5 dimerisation | unknown | 16 | 4:1 |

### 4.3 Optimisation Procedure

The parameters of the considered models are inferred using maximum likelihood based estimation. Specifically, given *n*_*t*_ measurements 𝒟 = *{*(*t*_*i*_, ***y***(*t*_*i*_)) |*i* = 1, …, *n*_*t*_*}*, the negative log likelihood function is defined as

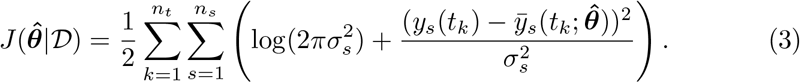

To prevent overfitting of the neural network, we add an L2 penalty to the objective function, yielding

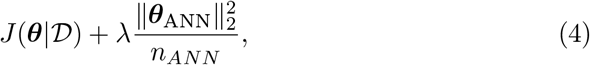

with regularisation parameter *λ* ≥0 and 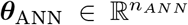. To minimize this objective function, we use a gradient-based optimisation approach: ADAM [22] for the first 500 epochs and BFGS [23] for the subsequent epochs (up to 3000). The optimised parameters are those that showed the minimal objective function value during optimisation on the validation set.

We implement a parameter transformation in order to work with optimisers that do not support parameter bounds or scales. Specifically, lower bounds 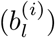 and upper bounds 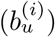 on 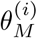, where *i* = 1, …, *n*_*M*_, 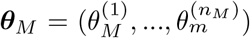, are realised with a scaled tanh function. Between the bounds, the tanh approximates a logarithmic function (see Fig. 8). Let *ρ*^(*i*)^ be the *i*th parameter of ***θ***_*M*_ on the transformed space. Then

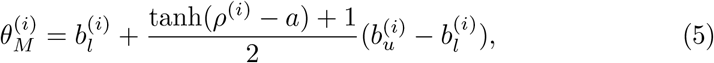

where *a* can be chosen such that - equivalent to the log-transform - 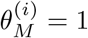 for *ρ*^(*i*)^ = 0.

**Fig. 8:**
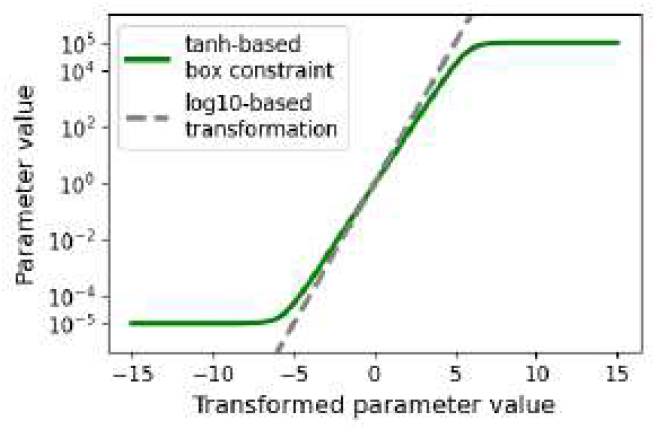
Tanh-based parameter transformation for *b*_*l*_ = 10^−5^ and *b*_*u*_ = 10^5^ in comparison to a log10 transformation.

With different start points of the mechanistic parameters and hyperparameter values according to Tab. 3, we run the optimisation several times. With each optimisation run, the initial mechanistic parameters values are set based on latin hypercube sampling within the parameter bounds. Equivalently, we sample the hyperparameters of the neural network and optimisation algorithm using latin hypercube sampling. The neural network parameters are sampled once per neural network architecture and mechanistic setting. Specifically, most neural network weights are initialized according to a Glorot uniform initialisation [33], the weights of the last layer and the bias are set to zero. If the normalisation of the neural network was activated, each input element to the first layer *x* is transformed according to

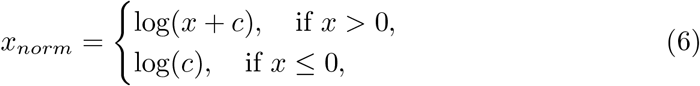

**Table 3:**
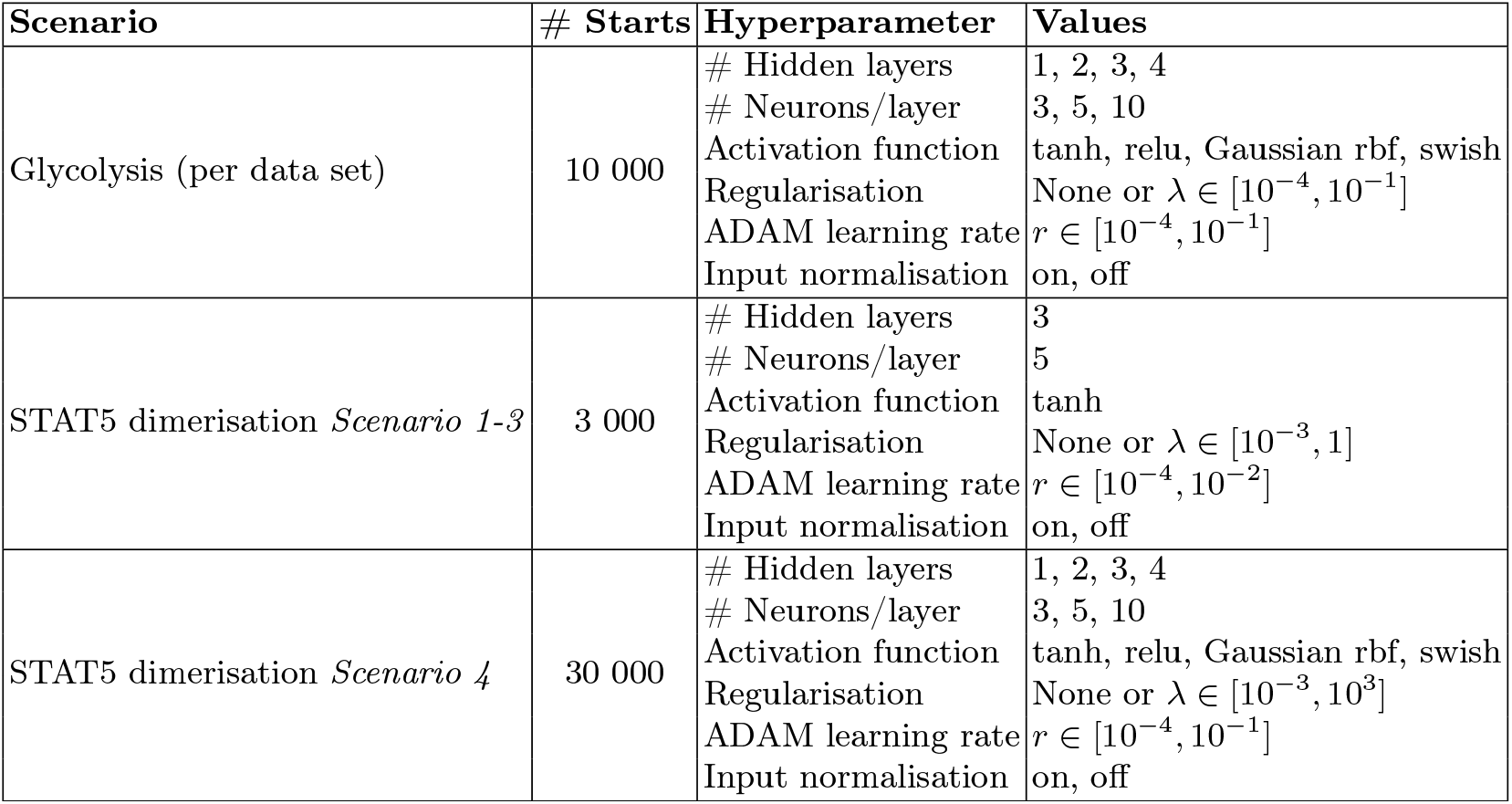
Overview of multi-start settings per problem and *Scenario*.

| Scenario | # Starts | Hyperparameter | Values |
| --- | --- | --- | --- |
| Glycolysis (per data set) | 10 000 | # Hidden layers<br># Neurons/layer<br>Activation function<br>Regularisation<br>ADAM learning rate<br>Input normalisation | 1, 2, 3, 4<br>3, 5, 10<br>tanh, relu, Gaussian rbf, swish<br>None or $\lambda \in [10^{-4}, 10^{-1}]$<br>$r \in [10^{-4}, 10^{-1}]$<br>on, off |
| STAT5 dimerisation *Scenario 1-3* | 3 000 | # Hidden layers<br># Neurons/layer<br>Activation function<br>Regularisation<br>ADAM learning rate<br>Input normalisation | 3<br>5<br>tanh<br>None or $\lambda \in [10^{-3}, 1]$<br>$r \in [10^{-4}, 10^{-2}]$<br>on, off |
| STAT5 dimerisation *Scenario 4* | 30 000 | # Hidden layers<br># Neurons/layer<br>Activation function<br>Regularisation<br>ADAM learning rate<br>Input normalisation | 1, 2, 3, 4<br>3, 5, 10<br>tanh, relu, Gaussian rbf, swish<br>None or $\lambda \in [10^{-3}, 10^3]$<br>$r \in [10^{-4}, 10^{-1}]$<br>on, off |

where we set *c* = 10^−20^ to ensure that the logarithm is defined for arbitrary values of *x*.

## 5 Data availability

Data sharing is not applicable to this article, as no datasets were generated or analysed during the current study.

## 6 Code availability

The underlying code for this study is available on GitHub and can be accessed via this link: https://github.com/m-philipps/ude_pipeline_systemsbio.

## 7 Acknowledgments

This work was supported by the Deutsche Forschungsgemeinschaft (DFG, German Research Foundation) under Germany’s Excellence Strategy (EXC 2047—390685813, EXC 2151—390873048) and the project ID 432325352 – SFB 1454, by the Federal Ministry of Education and Research (BMBF) under the CompLS program (GENImmune, grant no 031L0292F. and EMUNE, grant no 031L0293C) and the project INSIDe, grant no 031L0297A, and by the University of Bonn (via the Schlegel Professorship of J.H.). We thank Dr. Dilan Pathirana for his support in interpreting the analysis and preparing the manuscript. Optimisation was performed on the Unicorn and Marvin clusters at the University of Bonn.

## 8 Author contributions

J.H acquired funding and supervised the work. M.P. and N.S. conceptualised the study. M.P. and N.S. implemented the pipeline where N.S. implemented the julia-based optimisation workflow. M.P. evaluated the results and wrote the manuscript with help from N.S. and J.H. All authors reviewed the manuscript.

