## Supplement for "Universal differential equations for systems biology: Current state and open problems" (from another version of this preprint: version 1, 2024-12-05)

### 10 Supplementary

#### 10.1 Glycolysis

The estimated mechanistic parameters, their bounds and the scale used for optimisation is listed in Table 4.

| **Parameter name** | **Scale** | **Bounds** | **True value** | **UDE** |
|---|---|---|---|---|
| $J_0$ | log10 | 1e-1, 1e2 | 2.5 | Yes |
| $k_1$ | log10 | 1, 1e3 | 100 | Yes |
| $k_2$ | log10 | 1e-1, 1e2 | 6 | Yes |
| $k_3$ | log10 | 1e-1, 1e2 | 16 | Yes |
| $k_4$ | log10 | 1, 1e3 | 100 | Yes |
| $k_5$ | log10 | 1e-1, 1e2 | 1.28 | No |
| $k_6$ | log10 | 1e-1, 1e2 | 12 | Yes |
| $k_{ex}$ | log10 | 1e-1, 1e2 | 1.8 | Yes |
| kappa | log10 | 1e-1, 1e2 | 13 | Yes |
| $K_1$ | log10 | 1e-2, 1e1 | 0.52 | Yes |
| N | log10 | 1e-1, 1e2 | 1 | Yes |
| A | log10 | 1e-1, 1e2 | 4 | Yes |
| phi | log10 | 1e-2, 1e1 | 0.1 | Yes |
| sd $N_2$ (5% noise) | linear | 0, 0.025 | 0.00606935 | Yes |
| sd $N_2$ (10% noise) | linear | 0, 0.05 | 0.0121387 | Yes |
| sd $N_2$ (20% noise) | linear | 0, 0.1 | 0.0242774 | Yes |
| sd $N_2$ (35% noise) | linear | 0, 0.2 | 0.04248545 | Yes |
| sd $A_3$ (5% noise) | linear | 0, 0.4 | 0.0929227 | Yes |
| sd $A_3$ (10% noise) | linear | 0, 0.8 | 0.1858454 | Yes |
| sd $A_3$ (20% noise) | linear | 0, 1.6 | 0.3716908 | Yes |
| sd $A_3$ (35% noise) | linear | 0, 3.2 | 0.6504589 | Yes |

Table 4: Mechanistic parameters with their optimisation scale, bounds, true value, and whether it was optimised in the UDE as well.

A dense data set with altogether 61 data points was generated and subsampled to create sparse data sets with 46, 31, 16 and 8 data points in total. All data sets were noise-corrupted with additive normally-distributed noise and standard deviation of 5%, 10%, 20% or 35% of the nominal values. A dense ($\Delta t = 0.01$) and noise-free simulation in $t \in [1.5, 5]$ serves as the test set for all training data sets.

#### 10.2 STAT5 dimerisation

The estimated mechanistic parameters, their bounds and the scale used for optimisation is listed in Table 6. Unless indicated otherwise, all mechanistic parameters were used in all UDE *Settings*.

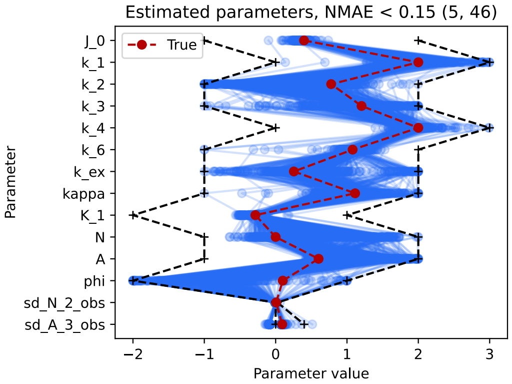


Fig. 9: Parameter vectors of the successful fits for the dataset with 5% noise and 46 data points.

| Reg. $\lambda$ | UDE ID | Symbolic regression | Complexity |
|---|---|---|---|
| 0 | 129253 | $0.03054 \cdot S_3 - 3.229$ | 5 |
| 1e-3 – 1e-2 | 129393 | $-107.3 \cdot (S_3 \cdot S_2) - 3.252$ | 7 |
| 1e-2 – 1e-1 | 126616 | $-8.778 \cdot {S_3}^2 - 3.054$ | 7 |
| 0.1 – 1 | 128044 | $-1.618 \cdot S_4 - 3.067$ | 7 |
| 1 – 10 | 124014 | $-8.057 \cdot S_3 - 2.527$ | 7 |

Table 5: Symbolic regression results for the glycolysis model, by regularisation strength $\lambda$. True solution: $-1.28 A_3$

.

| **Parameter name** | **Scale** | **Bounds** | **Reference value** | **Comment** |
|---|---|---|---|---|
| Epo_degradation_BaF3 | log10 | 1e-5, 1e5 | 0.026982514033029 | |
| k_exp_hetero | log10 | 1e-5, 1e5 | 1.00067973851508e-5 | |
| k_exp_homo | log10 | 1e-5, 1e5 | 0.006170228086381 | |
| k_imp_hetero | log10 | 1e-5, 1e5 | 0.0163679184468 | |
| k_imp_homo | log10 | 1e-5, 1e5 | 97749.3794024716 | |
| k_phos | log10 | 1e-5, 1e5 | 15766.5070195731 | |
| sd pSTAT5A_rel | log10 | 1e-5, 1e5 | 3.85261197844677 | |
| sd pSTAT5B_rel | log10 | 1e-5, 1e5 | 6.59147818673419 | |
| sd rSTAT5A_rel | log10 | 1e-5, 1e5 | 3.15271275648527 | |
| k_exp_aug_A | log10 | 1e-5, 1e5 | 0 | Only used in UDE *Scenario 4* |
| k_exp_aug_AB | log10 | 1e-5, 1e5 | 0 | |
| k_exp_aug_B | log10 | 1e-5, 1e5 | 0 | |

Table 6: STAT5 model mechanistic parameters with their optimisation scale, bounds, and reference value from the original publication [21].

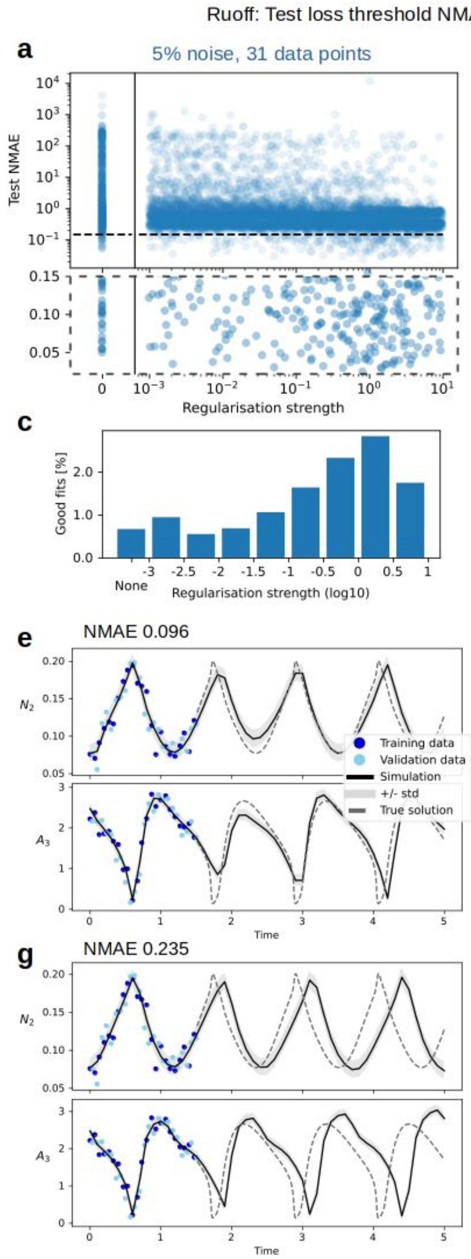


Fig. 10: Distribution of test losses and some fits above and below the NMAE test loss threshold 0.15. **a)** Scatter plot of test loss by regularisation strength for each optimised UDE, for the 5% noise and 31 data points; **b)** and for 10% noise and 46 data points. Additionally, the location of the UDEs from panels e-h are labeled. **e)** Predictions for a successful fit because the test loss fall below the threshold. **f-g)** predictions from failed models.

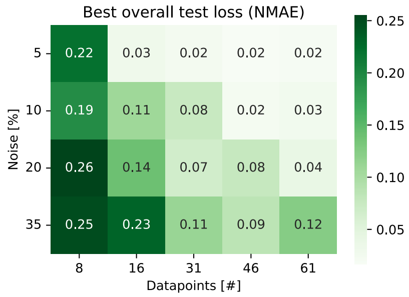


Fig. 11: The best test loss (NMAE) by data setting.

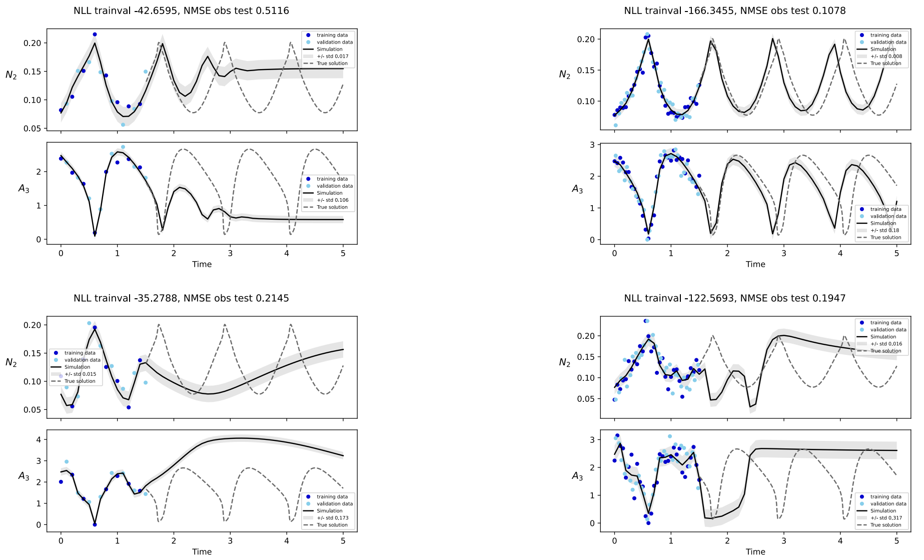


Fig. 12: Best fits from glycolysis model on datasets with 16 (left column) and 61 (right column) data points per observable; and lower noise (upper row, 10%) or higher noise (lower row, 20%).

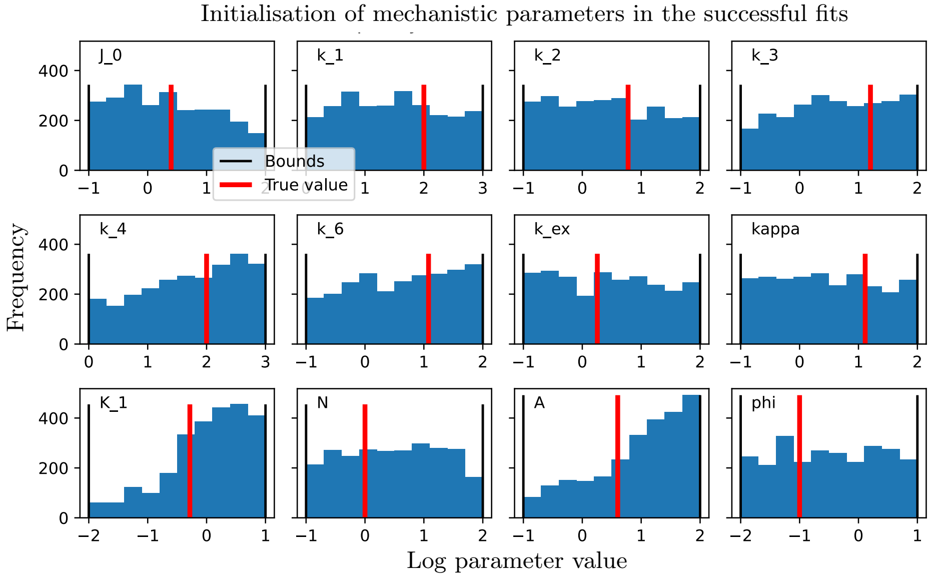


Fig. 13: Initial values for the mechanistic parameters, for all successfully fitted models (test loss $< 0.15$).

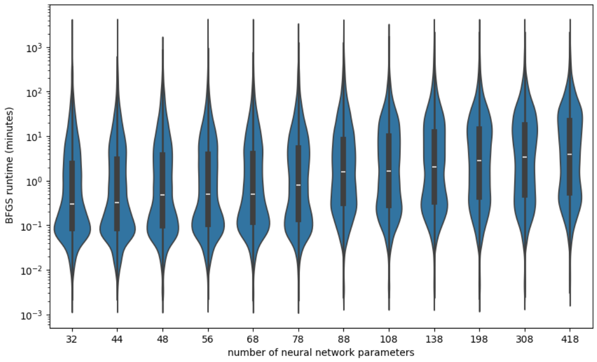


Fig. 14: Distribution of BFGS runtime by number of ANN parameters (weights + biases).

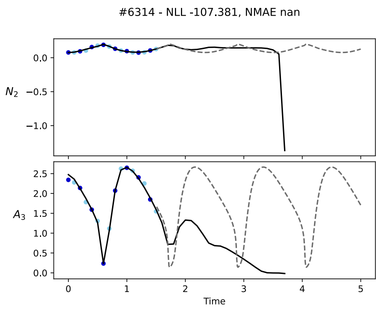


Fig. 15: UDE with best training loss that was trained on 16 data points with 5% noise without regularisation.

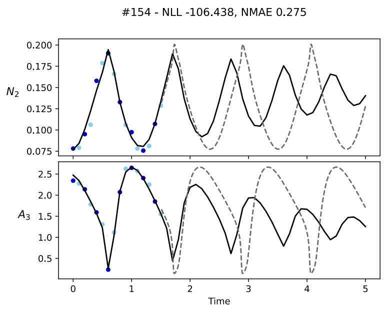


Fig. 16: UDE with best training loss that was trained on 16 data points with 5% noise with regularisation strength $1 < \lambda < 10$.

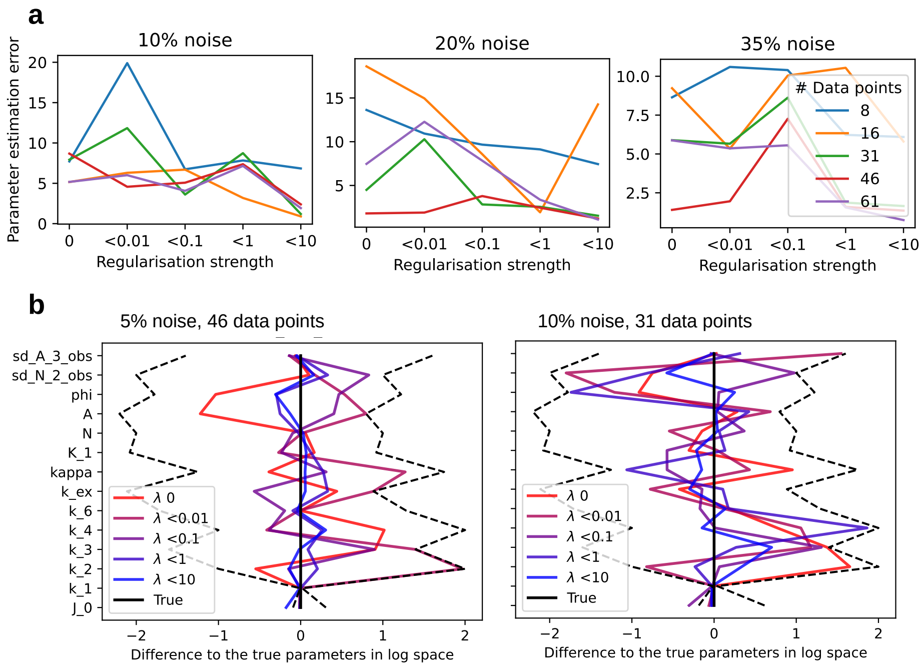


Fig. 17: **(a)** Parameter estimation error for the best UDEs by regularisation strength, shown for different data sets. **(b)** Parameter error per mechanistic parameter, i.e. difference to the true parameter, depending on the regularisation strength, show for two data sets.

| **Framework** | **Solver** | **Adaptive** | **Simulation Time** | **Simulation Success** |
|---|---|---|---|---|
| torchdiffeq | 8th-order Runge-Kutta of Dormand-Prince-Shampine | Yes | $> 12$h | No (timeout) |
| torchdiffeq | 5th-order Runge-Kutta of Dormand-Prince-Shampine | Yes | $> 12$h | No (timeout) |
| torchdiffeq | 3rd-order Runge-Kutta of Bogacki-Shampine | Yes | $> 12$h | No (timeout) |
| torchdiffeq | 2nd-order Runge-Kutta-Fehlberg | Yes | $> 12$h | No (timeout) |
| torchdiffeq | 2nd-order Runge-Kutta | Yes | $> 12$h | No (timeout) |
| SciML | KenCarp4 | Yes | 0.0003 s | Yes |
| SciML | Tsit5 | Yes | 0.303 s | Yes |

Table 7: Simulation results for all available torchdiffeq solvers and selected SciML solvers when simulating the original STAT5 dimerisation problem with true parameter values.

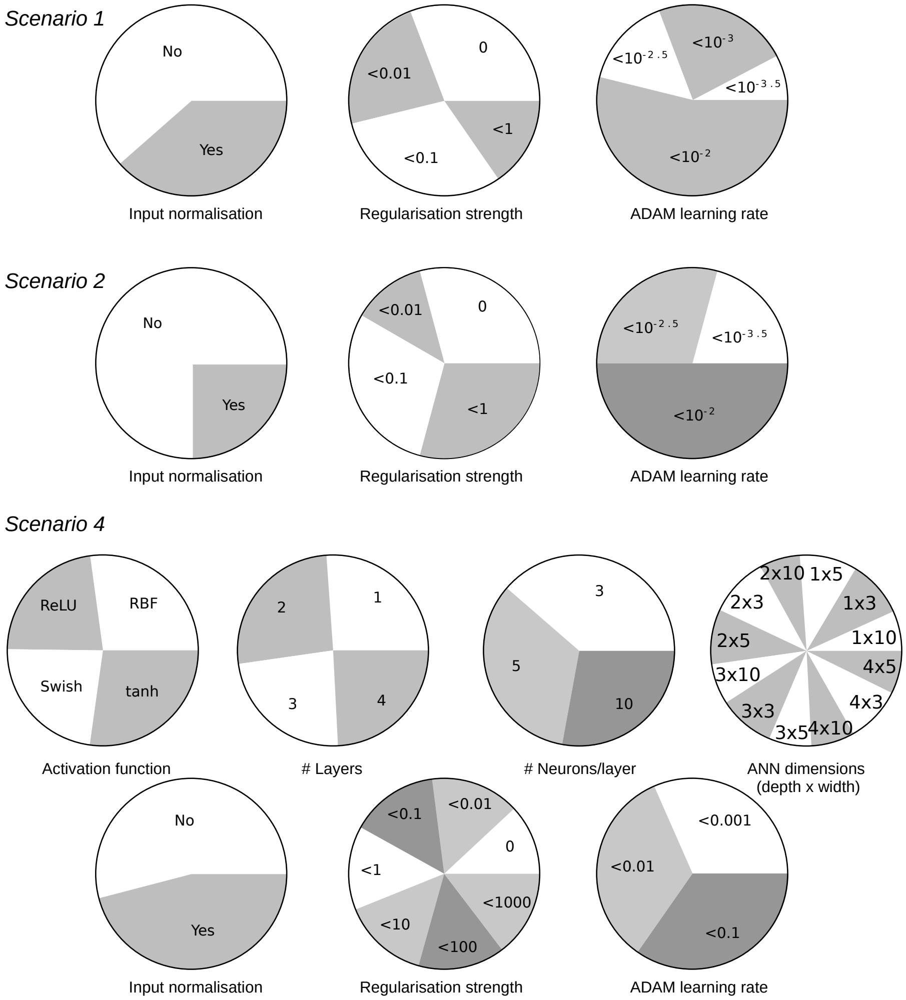


Fig. 18: For all successful fitted models of *Scenario 1, 2, 4*: Each pie represents one hyperparameter and the pie's fractions correspond to the number of successful fits with a specific setting. The continuously sampled hyperparameters (learning rate, regularisation strength) were binned, and the labels show the upper bound. Number of successful fitted models: 13 (*Scenario 2*), 24 (*Scenario 2*), 22719 (*Scenario 4*).
